# Effect of cow’s milk with different PUFA n-6: n-3 ratios on performance, serum lipid profile, and blood parameters of grower gilts

**DOI:** 10.1101/2021.10.04.463091

**Authors:** Leriana Garcia Reis, Thiago Henrique da Silva, Márcia Saladini Vieira Salles, André Furugen Cesar Andrade, Simone Maria Massami Kitamura Martins, Paula Lumy Takeuchi, Ana Maria Centola Vidal, Arlindo Saran Netto

**Author notes:** Corresponding author: Arlindo Saran Netto. /+55 19 35654039/+55 19 997270373.

## Abstract

The concern with human health has increased the interest in producing foods enriched with polyunsaturated fatty acids (PUFA), directly or naturally by inclusion in the animals’ diet. The positive effects such as antithrombotic, anti-inflammatory, and hypolipidemic have been observed in pigs and rats, used as human models for study. The present study evaluated the effect of cow’s milk with different lipid profiles on performance, serum fatty acid profile, biochemical analysis, and a complete blood count of gilts used as a human model. At 34 days, thirty gilts were equally distributed in three treatments. Experimental treatments were milk from cows without the oil supplementation (C), milk from cows fed an enriched diet with linseed oil (n-3), and milk from cows fed an enriched diet with soybean oil (n-6). Milk supplementation was performed until the age of 190 days, provided once in the morning. The n-3 and n-6 milk reduced the concentration of myristic acid in the blood and increased the leukocytes. Milk enriched with n-3 compared to n-6 reduced stearic acid. In conclusion, milk with a better PUFA profile can reduce saturated fatty acids in the blood and alter the concentration of cells in the defense system.

## INTRODUCTION

The consumption of cow’s milk and its derivatives in human nutrition has been reduced or even replaced by other beverages of vegetable origin due to the lipid profile [1] and also due to the occurrence of lactose intolerance and some proteins [2].

It is known that the composition of fatty acids in cow’s milk can be changed and is highly influenced by the nature of the forage consumed, supplementation with vegetable or oilseed oils, and even vitamin E and this change can be verified in a few hours [3,4]. Researchers have intensified studies in modifying the lipid profile in milk and meat [5,6]. More than 400 fatty acids compounded milk fat, and among them, saturated fatty acids (SFA) are in greater concentration. Excessive consumption of SFA has generated discussions because it is associated with an increased risk of developing several diseases, in addition to causing an increase in high-density lipoproteins (HDL) and low-density lipoproteins (LDL) cholesterol [7–9]. Research has been carried out to include different sources of unsaturated fatty acids (USFA), among them mainly polyunsaturated fatty acids (PUFA), in the diet of cows, focusing on improving the composition of milk fat and nutraceutical properties for human consumption [10]. High n-3 PUFA content and low n-6/n-3 PUFA ratio are more beneficial to human health, which n-3 and n-6 polyunsaturated fatty acids and their proportions are correlated with human diseases [11].

According to [12], the inclusion of canola oil in dairy cow diets can reduce milk SFA and improved omega-3. Results similar were obtained by [13] showed alteration on milk fatty acids profile after supplementing dairy cows with linseed and soybean oil, resulting in improved saturated/unsaturated fatty acid ratio and higher content of omega-3, and a more desirable omega-6/omega-3 ratio.

Milk enriched with PUFA showed a reduction in the concentration of LDL in the blood of rats [14], and in gilts supplemented with cow’s milk naturally enriched with n-3 and n-6 from 34 days of age until the end of lactation also showed a reduction in LDL for the n-3 and n-6 treatments and improve the performance of weaned piglets [15]. However, we did not find studies evaluating the effects of providing PUFA-enriched milk on the development of pigs, considering the pig as an animal model for humans. Several factors make some studies in humans unfeasible, such as the limitations of invasive procedures and the myriad of ethical issues, in addition to animal models being essential and relevant to support epidemiological studies [16]. In anatomy, neurobiology, cardiac vasculature, gastrointestinal tract, and genome, pigs are more human-like than mice and are currently considered adequate biomedical models [17]. Thus, the present study aimed to evaluate the supplementation of cow’s milk naturally enriched with n-3 and n-6 polyunsaturated fatty acids on swine’s health during the growing through growth performance, blood fatty acid profile, blood chemical variables, and complete blood counts.

## MATERIAL AND METHODS

All activities performed in this study were reviewed and approved by the Animal Care Committee of the School of Animal Science and Food Engineering, University of Sao Paulo (#4939070317).

### Animals, facilities, and treatment allocation

Thirty hybrid gilts (Landrace x Large White) were evaluated from 34 to 190 days old and were randomly allocated into three groups: control (**C**, n = 10), basal diet + milk from cows without the oil supplementation, omega-3 (**n-3**, n = 10), basal diet + milk from cows fed an enriched diet with linseed oil, and omega-6 (**n-6**, n = 10), basal diet + milk from cows fed an enriched diet with soybean oil.

At 34 days of age and initially weighing 9.59 ± 1.28 kg, the gilts were housed individually in pens (from 34 to 76 days, pre-initial and initial phase with 0.75 m^2^/animal; subsequently from 77 to 190 days old, grower-finisher phase until replacement phase, with 2.06 m^2^/gilt). The gilts were fed *ad libitum* twice a day (at 07:00 am and 1:00 pm) during the grower phase (130 days). After this period, the feed intake was adjusted according to [18]. The diets were based on corn and soybean feedstuffs and did not contain any oil as an ingredient (Table 1). All animals received *ad libitum* water.

**Table 1.** Composition of pre-initial, initial, growth and termination diets, specific to each physiological phase of swine females

| Item | Pre-initial | Initial | Grower | Finisher | Replacement | Gestation | Lactation |
| --- | --- | --- | --- | --- | --- | --- | --- |
| *Ingredients, g.kg<sup>-1</sup>* |  |  |  |  |  |  |  |
| Ground corn | 399.0 | 649.0 | 699.0 | 739.0 | 644.4 | 594.0 | 587.8 |
| Soybean meal | 200.0 | 300.0 | 280.0 | 240.0 | 240.8 | 140.0 | 265.1 |
| Wheat bran | - | - | - | - | 86.5 | 240.0 | - |
| UNIMIX<sup>a</sup> | 400.0 | 50.0 | 20.0 | 20.0 | 25.0 | 25.0 | 30.0 |
| L-lysine | - | - | - | - | - | - | 10.0 |
| DL-methionine | - | - | - | - | - | - | 2.0 |
| Sugar | - | - | - | - | - | - | 100.0 |
| Calcitic limestone | - | - | - | - | 2.3 | - | 4.1 |
| Mycofix | 1.0 | 1.0 | 1.0 | 1.0 | 1.0 | 1.0 | 1.0 |
| *Chemical composition* |  |  |  |  |  |  |  |
| Dry matter, % | 83.41 | 87.92 | 87.07 | 88.93 | 88.12 | 89.42 | 88.25 |
| Ashes, % | 6.69 | 5.43 | 4.80 | 3.70 | 4.54 | 5.23 | 5.62 |
| Crude energy, cal.g<sup>-1</sup> | 4347.0 | 4393.0 | 4419.0 | 4465.0 | 4411.5 | 4360.0 | 4376.0 |
| Ether extract, % | 4.14 | 1.71 | 2.41 | 1.71 | 1.37 | 1.40 | 1.56 |
| Crude fiber, % | 4.54 | 4.32 | 4.30 | 4.07 | 3.01 | 5.23 | 5.28 |
| Crude protein, % | 18.04 | 21.21 | 18.07 | 20.46 | 19.04 | 17.59 | 19.99 |
| Calcium, % | 0.90 | 0.80 | 0.63 | 0.51 | 0.82 | 0.81 | 0.94 |
| Phosphorus, % | 0.69 | 0.53 | 0.41 | 0.37 | 0.38 | 0.46 | 0.43 |

<sup>a</sup>The premix used in each phase was based on Rostagno et al. [18]

Daily milk supplementation followed the recommendation at approximately 5 mL/kg body weight according to the [19]. The milk was provided for pigs at 08:00 am, after individual feeding. Each gilt was supplemented with 200 mL (from 34 to 76 days), 300 mL (from 77 to 128 days), 400 mL (from 129 to 174 days), and 500 mL (175 to 190 days) of cow’s milk. The enriched milk provided to the gilts in this current study was obtained from the study carried out by [13] using Holstein cows. Briefly, the cows were supplemented or not with 2,5% (dry matter basis) of linseed or soybean oil, sources of n-3 and n-6, respectively. The effects of these vegetable oils’ inclusion on the lipid fraction profile of dairy cows’ milk are presented in Table 2.

**Table 2.**
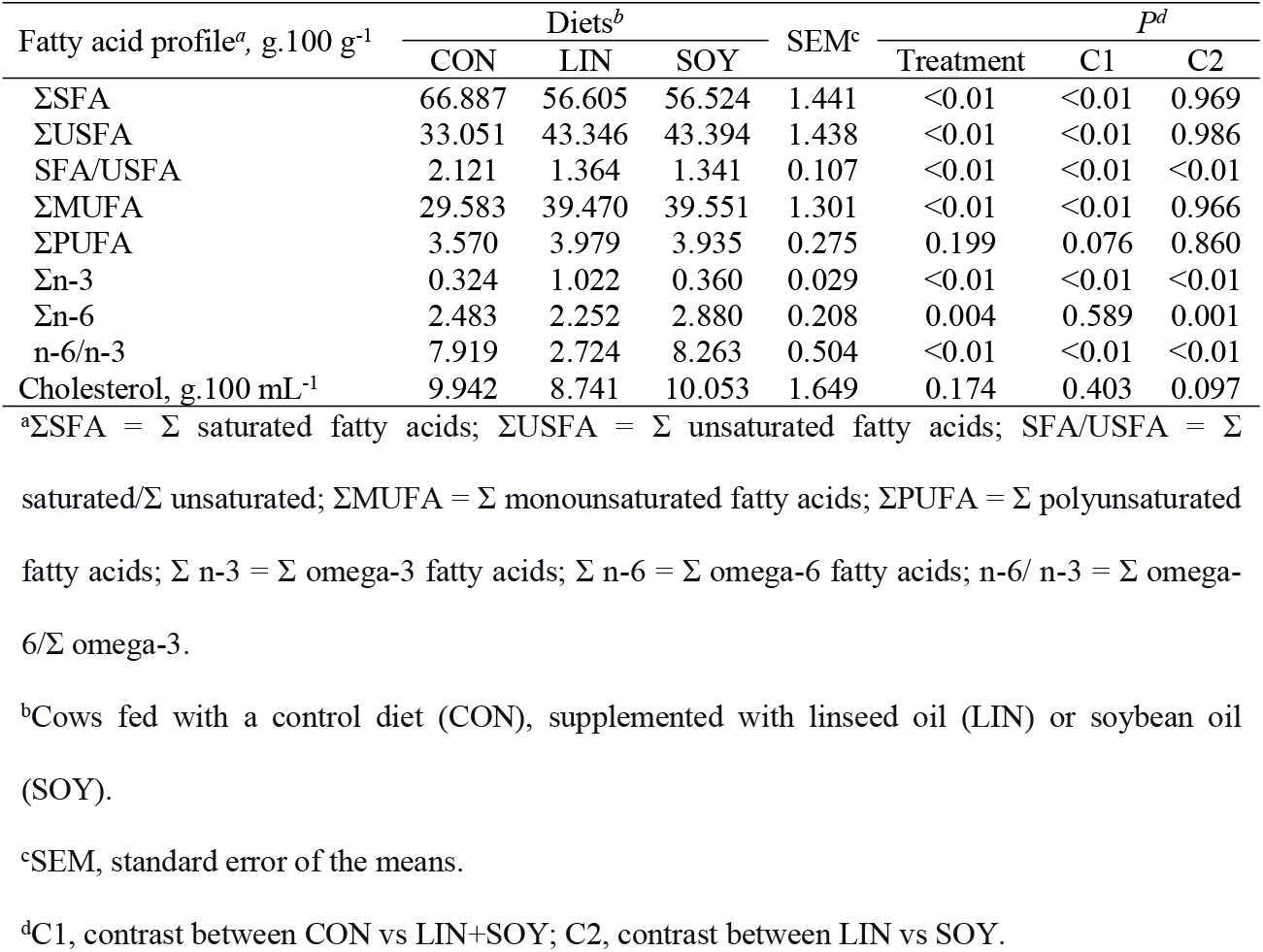
The lipid fraction of enriched cow’s milk

### Data collection

*Data of gilts were collected at* 34, 76, 112, 129, 155, and 190 days.

### Growth

Body weight (BW) was assessed using a digital scale (Toledo^®^, model MGR-3000, São Bernardo do Campo, SP), consisting of initial (from 34 to 76), grower (from 77 to 112), and finisher (from 113 to 190) phases. Additionally, average daily feed intake (ADFI), and from these measurements, the average daily gain (ADG) and the feed conversion ratio (FCR) were calculated.

### Blood sampling and analysis

Whole blood, serum, and plasma were collected by jugular venipuncture using a 1.2 × 40 mm collection needle (Becton Dickinson & Company, Juiz de Fora, MG, Brazil). Whole blood and serum were collected into 2 mL tubes with a clot activator, and plasma into 2 mL EDTA tubes (Vacuette^®^, Greiner Bio-One Brasil Produtos Médicos Hospitalares Ltda, Americana, SP, Brazil).

After collection, all tubes were immediately submitted to the laboratory (Diagnostic Laboratory for Clinical Analysis, Pirassununga, Brazil). The complete blood count was evaluated by Sysmex^®^ equipment (model XS-800i, Kobe, Japan). The serum sample was quantified by gas chromatography [20]. The plasma sample was analyzed for total cholesterol, HDL, total protein, and serum urea using enzyme kits (VIDA Biotecnologia^®^, Minas Gerais, Brazil). LDL and triacylglycerol were analyzed using enzyme kits (LABTEST^®^, Minas Gerais, Brazil). The very-low-density lipoprotein (VLDL) concentration was determined indirectly by the following equation [21]: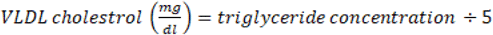. All analyses were according to the manufacturer’s instructions.

### Statistical Analyses

All statistical analyses were performed using SAS 9.4 (SAS Inst. Inc., Cary, NC). The data were analyzed in a completely randomized design, and the animal was considered an experimental unit. The animals were randomly distributed in 1 of 3 treatments. The normality of the residues was verified by the Shapiro-Wilk test (UNIVARIATE procedure of SAS), and information with student residues greater than +3 or less than −3 was excluded from the analyzes. The Levene test compared the homogeneity of the variances. Variables with a continuous distribution such as body weight, average daily gain, average daily feed intake, feed conversion rate, complete blood counts, biochemical parameters, and fatty acids profile of blood were analyzed using the MIXED procedure of SAS. When the time factor was present, repeated measures models were fitted by multiple mixed linear models using the MIXED procedure of SAS. The model included: the fixed effects of treatment, time and their interaction, and the random effect of the animal. The experimental unit was the gilt. The SLICE option using the LSMEANS/PDIFF command was used to explore the interactions between treatment and day of data collection. The Kenward-Roger method was used to correct the degree of freedom of the denominator for the F test. The covariance structure was determined based on the lowest Akaike information criteria value. The treatments were analyzed using two orthogonal contrasts: contrast 1 (C1) control group vs. n-3 + n-6 groups, and contrast 2 (C2) n-3 group vs. n-6 group. Significance level was set at P < 0.05 and tendency towards significance at 0.05 < P < 0.10.

## RESULTS

All gilts consumed their supplementation successfully so that the milk remains were not significant.

### Gilts’ performance

There was no difference in the treatment x time interaction on performance. Females supplemented with n-3 and n-6 had similar growth performance between themselves, and in comparison, to those of the control groups (see S1 Fig). Only time effect was observed for all these variables (P < 0.001, Table 3).

**Table 3.** Growth performance of gilts from 34 to 190 days of age, for treatments C, n-3 and n-6

| Item<sup>a</sup> | Treatments<sup>b</sup> |  |  | SEM<sup>c</sup> | P-value |  |  |  |  |
| --- | --- | --- | --- | --- | --- | --- | --- | --- | --- |
|  | Control | n-3 | n-6 |  | Treatment | Time | Treat*Time | C1<sup>d</sup> | C2<sup>d</sup> |
| BW, kg | 67.570 | 67.540 | 66.750 | 2.654 | 0.942 | <0.0001 | 0.929 | 0.947 | 0.915 |
| ADG, kg.dia<sup>-1</sup> | 0.754 | 0.750 | 0.734 | 0.044 | 0.864 | <0.0001 | 0.967 | 0.760 | 0.729 |
| ADFI, kg.dia<sup>-1</sup> | 2.348 | 2.356 | 2.328 | 0.046 | 0.846 | <0.0001 | 0.949 | 0.951 | 0.812 |
| FC ratio, kg.kg<sup>-1</sup> | 3.138 | 3.147 | 2.990 | 0.063 | 0.870 | <0.0001 | 0.948 | 0.598 | 0.323 |

<sup>a</sup>BW: body weight; ADG: average daily gain; ADFI: average daily feed intake; FC ratio: feed:gain ratio.
<sup>b</sup>Sows fed with a Control milk (Control), supplemented with cow's milk enriched with n-3 or n-6.
<sup>c</sup>SEM, standard error of the mean.
<sup>d</sup>C1, contrast Control vs n-3+n-6; C2, contrast n-3 vs n-6.

S1 Fig. Growth performance of females from 34 to 190 days old

### Fatty Acids Profile in Gilts Serum

At 77 days old, the control group had greater myristic acid concentration than those that received enriched cow’s milk (n-3 and n-6 P = 0.062, Fig 1 and S2 fig). At 132 days old, the n-3 gilts had an increased γ-linolenic acid concentration than other groups (P = 0.079, Fig 2). The control gilts group had raised myristic acid concentration compared to the females of other groups (P = 0.031, Table 4). Besides, the n-3 gilts group had a small SFA concentration (P = 0.090) and ratio SFA: USFA, (P = 0.097) and greater USFA than those of control and n-6 groups (P = 0.090, Table 4). Also, the n-3 female groups had lower stearic acid concentration (P = 0.002, Table 4 and S2 fig) and greater oleic acid concentration than those of the n-6 group (P = 0.081, Table 4). The other fatty acids were similar among groups (P > 0.05, Table 4).

**Fig 1.**
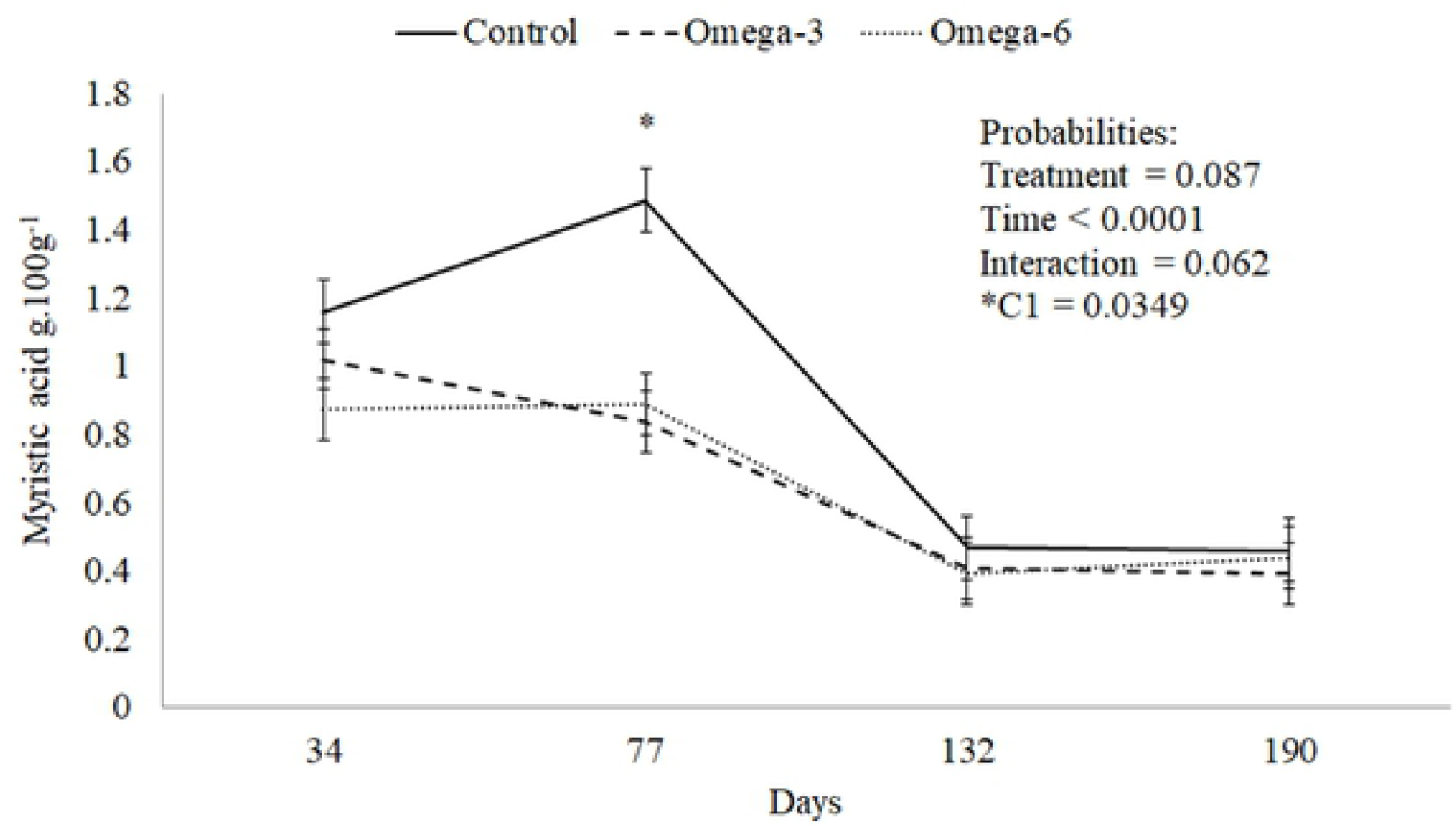
Tendency to increase myristic acid for Control at 77 days of age of gilts

**Fig 2.**
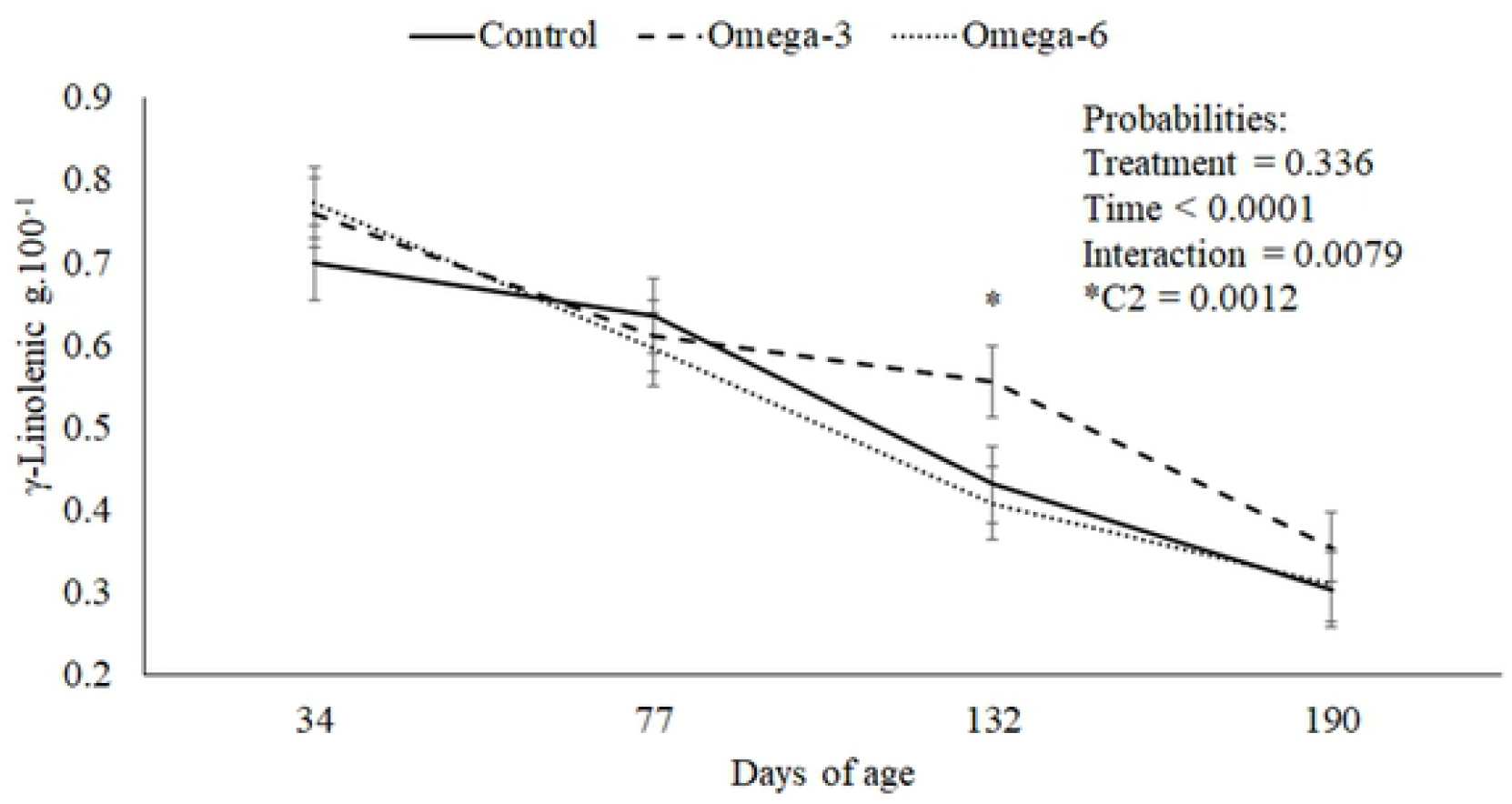
γ-linolenic tends to have a higher concentration in n-3 treatment at 132 days of age of gilts

**Table 4.** Serum quantitative fatty acid profile for treatments Control, n-3 and n-6 of females supplemented from 34 to 190 days of age

| Fatty acids<sup>a</sup>, µg.ml<sup>-1</sup> | Treatments<sup>b</sup> |  |  | SEM<sup>c</sup> | P-value |  |  |  |  |
| --- | --- | --- | --- | --- | --- | --- | --- | --- | --- |
|  | Control | n-3 | n-6 |  | Treatment | Time | Treat*Time | C1<sup>d</sup> | C2<sup>d</sup> |
| Myristic, C14:0 | 0.869 | 0.669 | 0.653 | 0.070 | 0.087 | <0.0001 | 0.062 | 0.031 | 0.875 |
| Palmitic, C16:0 | 17.730 | 17.800 | 17.490 | 0.713 | 0.950 | 0.0465 | 0.541 | 0.925 | 0.766 |
| Stearic, C18:0 | 12.430 | 11.830 | 13.560 | 0.304 | 0.006 | <0.0001 | 0.408 | 0.497 | 0.002 |
| Elaidic, C18:1t n-9 | 0.382 | 0.293 | 0.379 | 0.079 | 0.686 | 0.0024 | 0.174 | 0.652 | 0.470 |
| Palmitoleic, C16:1c9 | 1.136 | 1.167 | 1.117 | 0.097 | 0.936 | 0.0016 | 0.937 | 0.961 | 0.721 |
| Oleic, C18:1c9 | 23.330 | 24.050 | 22.130 | 0.714 | 0.200 | 0.0028 | 0.684 | 0.790 | 0.081 |
| Linoleic, C18:2 n-6 | 28.390 | 29.770 | 28.670 | 0.545 | 0.225 | 0.0001 | 0.708 | 0.242 | 0.190 |
| Dihomo-γ-linolenic, C20:3 n-6 | 0.695 | 0.688 | 0.660 | 0.045 | 0.852 | <0.0001 | 0.637 | 0.714 | 0.679 |
| ARA, C20:4 n-6 | 11.240 | 11.000 | 12.410 | 0.589 | 0.239 | <0.0001 | 0.587 | 0.534 | 0.121 |
| γ-Linolenic, C18:3 n-6 | 0.506 | 0.567 | 0.531 | 0.028 | 0.336 | <0.0001 | 0.079 | 0.232 | 0.381 |
| α-Linolenic, C18:3 n-3 | 0.563 | 0.618 | 0.588 | 0.026 | 0.361 | <0.0001 | 0.326 | 0.232 | 0.431 |
| EPA, C20:5 n-3 | 0.219 | 0.343 | 0.461 | 0.096 | 0.239 | 0.1157 | 0.280 | 0.143 | 0.397 |
| DHA, C22:6 n-3 | 1.203 | 1.184 | 1.371 | 0.112 | 0.459 | <0.0001 | 0.300 | 0.617 | 0.260 |
| Saturated | 31.410 | 30.370 | 31.150 | 0.339 | 0.090 | <0.0001 | 0.450 | 0.124 | 0.106 |
| Unsaturated | 68.590 | 69.630 | 68.850 | 0.323 | 0.090 | <0.0001 | 0.450 | 0.124 | 0.106 |
| SFA/USFA | 0.458 | 0.438 | 0.453 | 0.006 | 0.097 | <0.0001 | 0.472 | 0.132 | 0.110 |
| Monounsaturated | 25.240 | 25.430 | 24.040 | 0.675 | 0.312 | 0.0005 | 0.234 | 0.547 | 0.164 |
| Polyunsaturated | 43.340 | 44.200 | 44.880 | 0.817 | 0.438 | 0.2194 | 0.229 | 0.255 | 0.563 |
| Total n-3 | 2.002 | 2.154 | 2.384 | 0.142 | 0.197 | <0.0001 | 0.296 | 0.152 | 0.262 |
| Total n-6 | 41.350 | 42.050 | 42.480 | 0.775 | 0.596 | 0.0247 | 0.458 | 0.354 | 0.694 |
| n-6/n-3 | 27.271 | 26.347 | 25.528 | 1.112 | 0.559 | <0.0001 | 0.459 | 0.341 | 0.612 |
| ARA/EPA | 59.854 | 57.438 | 46.762 | 6.832 | 0.379 | 0.0043 | 0.174 | 0.368 | 0.285 |

<sup>a</sup>EPA: eicosapentaenoic acid; DHA: docosahexaenoic acid; ARA: arachidonic acid; USFA/SFA: unsaturated/saturated fatty acids; Total n-3: sum of n-3 fatty acids; Total n-6: sum of n-6 fatty acids; n-6/n-3, ratio of total n-6 to total n-3 fatty acids; ARA/EPA: ratio of arachidonic acid to eicosapentaenoic acid.
<sup>b</sup>Sows fed with a Control milk (Control), supplemented with cow's milk enriched with n-3 or n-6.
<sup>c</sup>SEM, standard error of the mean.
<sup>d</sup>C1, contrast Control vs n-3+n-6; C2, contrast n-3 vs n-6.

S2 Fig. Myristic and stearic acid of females from 34 to 190 days old

### Biochemical Parameters

At 77 days old, the n-6 gilts group had greater LDL than those of other groups (P = 0.083, Fig 3). Besides, these females increased HDL compared to the n-3 group (P = 0.063, Table 5). The other biochemical parameters were similar among females of the control group and those supplemented with enriched cow’s milk (P > 0.05).

**Fig 3.**
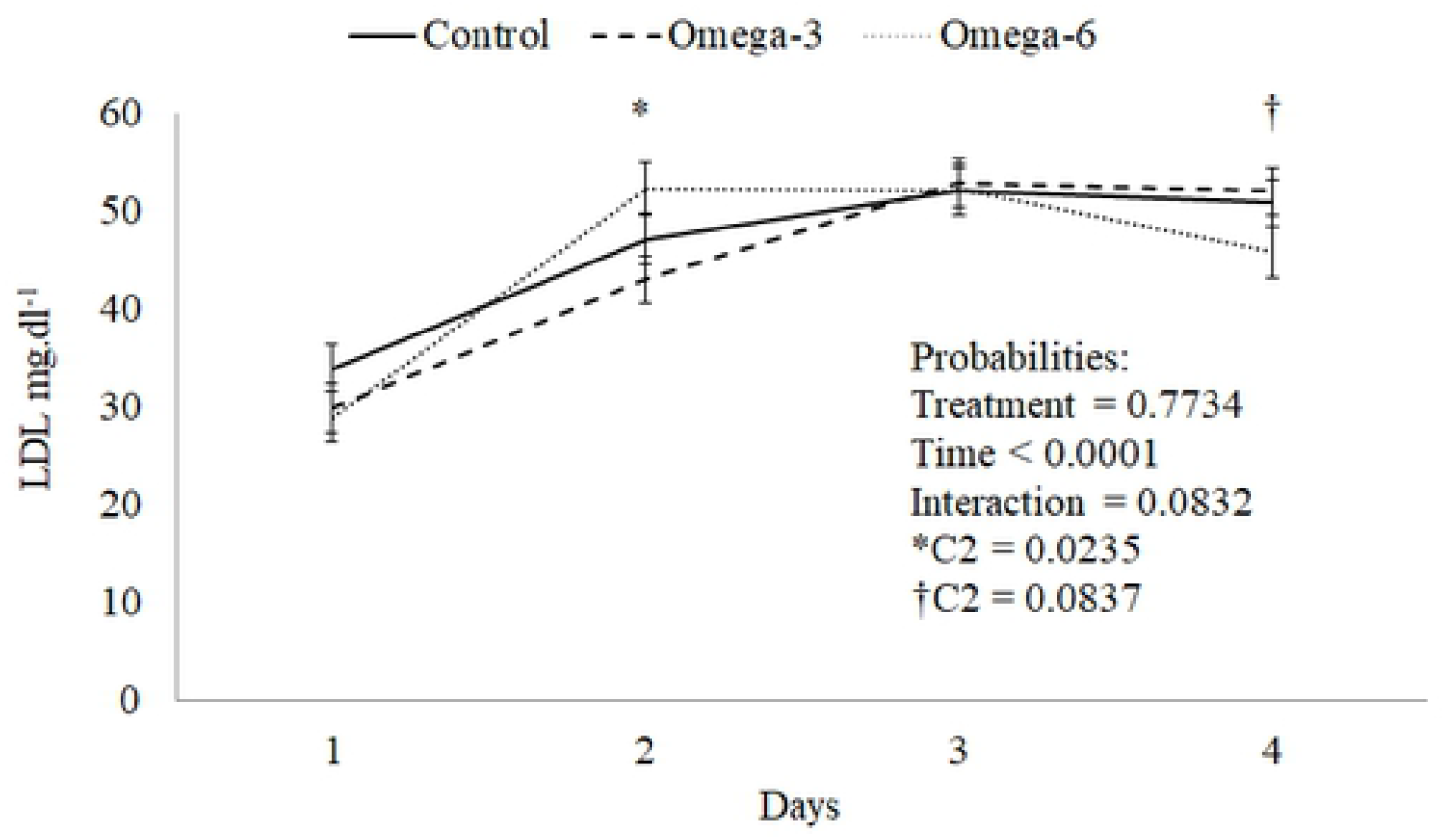
Behavior of the LDL curve at 77 days of gilts

**Table 5.** Biochemical parameters of the serum of gilts from 34 to 190 days of age

| Biochemical parameters | Treatments<sup>a</sup> |  |  | SEM<sup>b</sup> | P-value |  |  |  |  |
| --- | --- | --- | --- | --- | --- | --- | --- | --- | --- |
|  | Control | n-3 | n-6 |  | Treatment | Time | Treat*Time | C1<sup>c</sup> | C2<sup>c</sup> |
| Urea, mg.dL<sup>-1</sup> | 23.590 | 24.370 | 24.380 | 1.060 | 0.833 | <0.0001 | 0.851 | 0.722 | 0.978 |
| Total protein, g.dL<sup>-1</sup> | 5.760 | 5.720 | 5.810 | 0.070 | 0.661 | <0.0001 | 0.332 | 0.959 | 0.634 |
| Total cholesterol, mg.dL<sup>-1</sup> | 79.180 | 77.110 | 79.350 | 2.290 | 0.747 | <0.0001 | 0.181 | 0.687 | 0.490 |
| Glucose, mg.dL<sup>-1</sup> | 90.330 | 85.510 | 86.490 | 1.620 | 0.101 | <0.0001 | 0.198 | 0.133 | 0.768 |
| Triglycerides, mg.dL<sup>-1</sup> | 35.890 | 37.300 | 32.490 | 2.860 | 0.501 | <0.0001 | 0.517 | 0.751 | 0.275 |
| HDL, mg.dL<sup>-1</sup> | 25.990 | 25.050 | 28.020 | 1.030 | 0.145 | <0.0001 | 0.812 | 0.730 | 0.063 |
| LDL, mg.dL<sup>-1</sup> | 46.020 | 44.510 | 44.910 | 1.550 | 0.773 | <0.0001 | 0.083 | 0.540 | 0.861 |
| VLDL, mg.dL<sup>-1</sup> | 7.200 | 7.520 | 6.690 | 0.560 | 0.595 | <0.0001 | 0.782 | 0.857 | 0.337 |

<sup>a</sup>Sows, fed with a Control milk (Control), supplemented with cow's milk enriched with n-3 or n-6.
<sup>b</sup>SEM, standard error of the mean.
<sup>c</sup>C1, contrast Control vs n-3+n-6; C2, contrast n-3 vs n-6.

### Hemogram and leukogram

Control gilts had reduced red blood cells (RBC), platelets, and lymphocytes compared to those that received enriched cow’s milk (P < 0.10, Table 6). Besides, these females had lower leukocytes (P = 0.049, Table 6) and higher monocytes (P = 0.041, Table 6) than the gilts of the omega group. These females also had increased mean corpuscular hemoglobin (MCH) compared to those of the omega group (P = 0.070, Table 6). The other counts were not affected by milk supplementation (P > 0.05, Table 6).

**Table 6.**
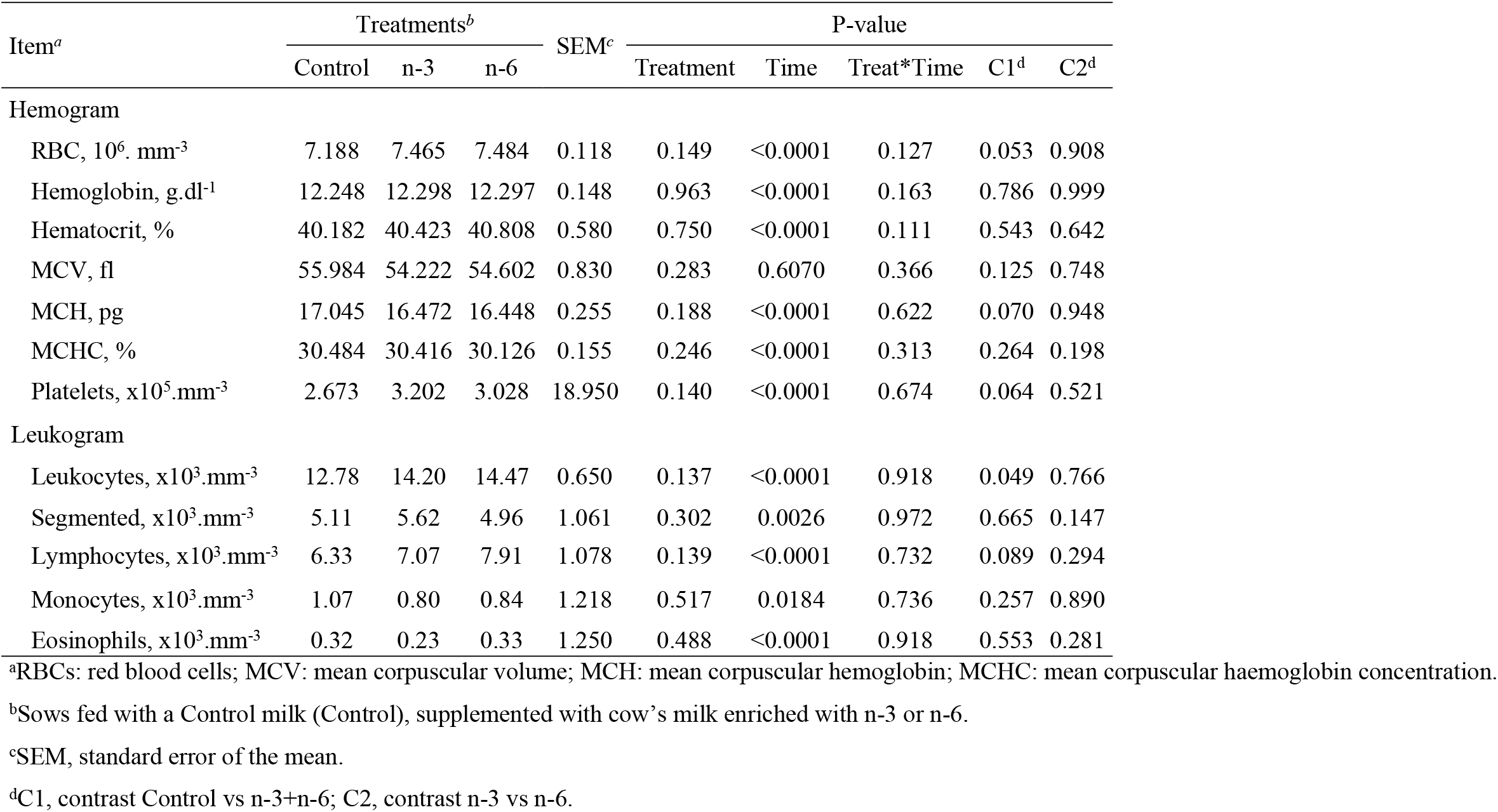
Hemogram and leukogram of the control, n-3, and n-6 groups of gilts supplemented from 34 to 190 days old

| Item<sup>a</sup> | Treatments<sup>b</sup> |  |  | SEM<sup>c</sup> | P-value |  |  |  |  |
| --- | --- | --- | --- | --- | --- | --- | --- | --- | --- |
|  | Control | n-3 | n-6 |  | Treatment | Time | Treat*Time | C1<sup>d</sup> | C2<sup>d</sup> |
| Hemogram |  |  |  |  |  |  |  |  |  |
| RBC, 10<sup>6</sup>. mm<sup>-3</sup> | 7.188 | 7.465 | 7.484 | 0.118 | 0.149 | <0.0001 | 0.127 | 0.053 | 0.908 |
| Hemoglobin, g.dl<sup>-1</sup> | 12.248 | 12.298 | 12.297 | 0.148 | 0.963 | <0.0001 | 0.163 | 0.786 | 0.999 |
| Hematocrit, % | 40.182 | 40.423 | 40.808 | 0.580 | 0.750 | <0.0001 | 0.111 | 0.543 | 0.642 |
| MCV, fl | 55.984 | 54.222 | 54.602 | 0.830 | 0.283 | 0.6070 | 0.366 | 0.125 | 0.748 |
| MCH, pg | 17.045 | 16.472 | 16.448 | 0.255 | 0.188 | <0.0001 | 0.622 | 0.070 | 0.948 |
| MCHC, % | 30.484 | 30.416 | 30.126 | 0.155 | 0.246 | <0.0001 | 0.313 | 0.264 | 0.198 |
| Platelets, x10<sup>5</sup>.mm<sup>-3</sup> | 2.673 | 3.202 | 3.028 | 18.950 | 0.140 | <0.0001 | 0.674 | 0.064 | 0.521 |
| Leukogram |  |  |  |  |  |  |  |  |  |
| Leukocytes, x10<sup>3</sup>.mm<sup>-3</sup> | 12.78 | 14.20 | 14.47 | 0.650 | 0.137 | <0.0001 | 0.918 | 0.049 | 0.766 |
| Segmented, x10<sup>3</sup>.mm<sup>-3</sup> | 5.11 | 5.62 | 4.96 | 1.061 | 0.302 | 0.0026 | 0.972 | 0.665 | 0.147 |
| Lymphocytes, x10<sup>3</sup>.mm<sup>-3</sup> | 6.33 | 7.07 | 7.91 | 1.078 | 0.139 | <0.0001 | 0.732 | 0.089 | 0.294 |
| Monocytes, x10<sup>3</sup>.mm<sup>-3</sup> | 1.07 | 0.80 | 0.84 | 1.218 | 0.517 | 0.0184 | 0.736 | 0.257 | 0.890 |
| Eosinophils, x10<sup>3</sup>.mm<sup>-3</sup> | 0.32 | 0.23 | 0.33 | 1.250 | 0.488 | <0.0001 | 0.918 | 0.553 | 0.281 |

<sup>a</sup>RBCs: red blood cells; MCV: mean corpuscular volume; MCH: mean corpuscular hemoglobin; MCHC: mean corpuscular haemoglobin concentration.
<sup>b</sup>Sows fed with a Control milk (Control), supplemented with cow's milk enriched with n-3 or n-6.
<sup>c</sup>SEM, standard error of the mean.
<sup>d</sup>C1, contrast Control vs n-3+n-6; C2, contrast n-3 vs n-6.

## DISCUSSION

Polyunsaturated fatty acids are essential for normal physiological function and the health of humans and domestic animals. Considering the opposing effects of n−3 and n−6 PUFA, the proportion of these PUFA, especially dietary linoleic acid and α-linolenic acid, may regulate diverse sets of homeostatic processes by themselves or by locally acting bioactive signaling lipids called eicosanoids derived from arachidonic acid, eicosapentaenoic acid, and docosahexaenoic acid [22].

Through the consumption of PUFA by diet, it is possible to reduce plasma levels of LDL and triacylglycerols. Therefore, SFA has a high potential to increase LDL and long-chain polyunsaturated fatty acids n-6 and n-3 to reduce it [23]. The supplementation of gilts with cow’s milk enriched naturally with PUFA from 34 to 190 days old led to changes in fatty acid profile and leukogram, as well as a trend was observed for increase HDL and reduce LDL, and hemogram; however, the growth performance had been similar among the females’ groups.

Liu and Kim [24] did not report any change in the growth performance of crossbred pigs, fed diets in different n-6: n-3 ratios (15.17, 9.68, and 4.87, high, medium, and lower, respectively). Nevertheless, in blood lipids, a reduction in total cholesterol, LDL, and triglycerides were observed for the lower ratio n-6:n-3 PUFA compared to animals in the other groups. Similarly, Li et al. [25] reported based on a meta-analysis in humans that low-ratio n-6/n-3 PUFA reduced triglyceride concentration and increased HDL concentration (P < 0.05), and the beneficial effects of low-ratio n-6/n-3 PUFA on blood lipids were enhanced with time. Besides these, researchers also verified that n-3 PUFA derived from plants significantly reduced total cholesterol and LDL concentrations, and n-3 PUFA derived from eicosapentaenoic acid (EPA) and DHA significantly reduced triglyceride concentration and increased HDL concentration.

In the present study, different n-6:n-3 PUFA ratios were observed in cow’s milk enriched with n-3 and n-6 sources (2.72, 7.92 and 8.26, n-3, group control, and n-6, respectively), but in female serum, the ratio was similar (16.35, 27.27 and 25.53, n-3, control group, and n-6, respectively) after 22 weeks of supplementation. Growth performance results were similar to those reported by Liu and Kim [24]. On the other hand, HDL trended an increase in the n-6 group compared with n-3, and total cholesterol was similar among the group. Considering the overall period, the LDL was numerically lower in animals from the n-3 and n-6 groups, but a trend towards the difference between the PUFA groups was observed at 77 and 190 days old. An increase in LDL concentration from 34 to 77 days was verified in three groups, but from that moment on, the animals in the n-6 group had a reduction while the other groups remained elevated. Several factors may influence these results, such as the presence or absence of lipoprotein disorders, the relation between VLDL and total cholesterol, the dose of oil administered, varying effects of EPA vs. DHA, the source of n-3, the type of individual the study is being conducted on (normolipidemic or hypercholesterolemic), and the period of supplementation [26,27].

The importance of lipids consumed in the diet on the effects on human health has been widely discussed, as they influence the lipid composition of blood, colostrum, milk, tissues, and it is even possible to observe the manipulation of gene expression and immunity [28–30]. Myristic and stearic acids are the most abundant saturated fatty acids in mammals [31]. Myristic acid has from 8 to 12% of total fat milk, and a more substantial intake of this lipid raises LDL concentrations [32].

Cow’s milk from the study by [13] was supplemented for gilts in this current study, and this milk had a lower concentration of myristic acid than those without the oil supplementation (7.59, 7.98, and 10.36, soybean, linseed, and control, respectively, P < 0.05), as well as reduction of SFA (56.52, 56.69 and 66.89, soybean, linseed, and control, respectively, P < 0.05). This fact may have directly influenced the serum fatty acid profile of gilts of n-3 and n6 groups. Besides, myristic acid may be produced through *de novo* lipogenesis, but it represents a small fraction [33]. Thus, we believe that the alteration found in the blood myristic acid may have come directly from the lipids present in milk through the transfer from the small intestine by diffusion or facilitated transport.

Higher concentration of stearic acid was observed in milk from cows that received soybean oil and linseed compared to control (14.50, 14.94, and 11.28, soybean, linseed, and control, respectively, P < 0.05), and no differences between the PUFA-supplemented groups were reported by [13]. The higher concentration of stearic acid found in the serum of gilts from the n-6 group may have been originated from *de novo* lipogenesis due to the higher elongation activity of palmitic acid by fatty acid elongase. Furthermore, PUFA is known to repress the stearoyl-CoA desaturase-1 (SCD-1) gene expression [34] that it is a lipogenic enzyme responsible for the biosynthesis of oleic acid (18:1) by desaturating stearic acid (18:0) [35–37]. The length, structure, and number of double bonds in the fatty acid chains also seem to influence the activity of SCD, according to [38]. The linoleic acid and EPA can directly regulate SCD activity at the level of transcription via sterol regulatory element-binding protein 1c (SREBP-1c), but the same does not occur with α-linolenic acid (ALA) [39]. Therefore, the difference in stearic acid found between n-3 and n-6 groups may have been influenced by all these factors, since cow’s milk enriched with soy oil had a higher concentration of linoleic acid (2.13, 2.62 and 2.03, control, soybean, and linseed, respectively [13].

In the present study, control gilts tended to have a reduced RBC, platelets, and lymphocytes and enhanced MCH compared to those that received enriched cow’s milk. These females also had lower leukocytes and higher monocytes than the gilts of the PUFA group.

Inflammation is a positive state for the body, as it denotes an immediate response to injury or infection. Thus, several cells are moved towards the affected site, including leukocytes, which can be modulated by fatty acids. This is because leukocytes have an important role in inflammation and infection, through the synthesis of important cytokines, some with more inflammatory functions, arising from PUFA n-6, and others with more anti-inflammatory functions, mainly from PUFA n-3 [40,41]. In this case, the n-6 PUFA is a substrate for the synthesis of eicosanoids, such as prostaglandins (PGs), thromboxanes (TXs), leukotrienes (LTs), and hydroxyeicosatetraenoic acids (HETEs). Among the series 4 LTs are: LTA_4_, B_4_, C_4_, D_4_ and E_4_, and LTB_4_ is a potent chemotactic agent for leukocytes [40].

Thus, it is possible that the PUFA-enriched milk, treatments n-3 and n-6 in this study, have stimulated this higher concentration of leukocytes due to the performance they can perform in the body in terms of cytokine production, cell proliferation, adhesion molecules, and cell death.

The consumption of PUFA can cause different effects on the cells that make up the blood plasma. According to Xia et al. [42], dietary n-3 polyunsaturated fatty acids (fish oil) in mice alters the hematopoietic microenvironment and induces the expression of matrix metalloproteinase 12 (MMP12) in stromal cells, providing new insights into diet-mediated regulation of hematopoiesis. Similar results were found by [43], in which oral administration of PUFA in mice increases hematopoiesis in them by generating active metabolites, which activate the Wnt and CXC Chemokine Receptor Type 4 (CXCR4) pathways and promote thrombopoiesis through the Notch proteins (Notch1) pathway. Although, how the dietary fatty acids can influence the hematopoiesis remains yet largely unclear. Unlike the aforementioned results, except for the segmented ones, the other variables related to hematopoiesis had similar results between groups n-3 and n-6, without the effect of n-3 PUFA in our study.

The increased intake of EPA and DHA replaces the arachidonic acid in the cell membranes of erythrocytes, platelets, neutrophils, monocytes, and liver cells [44] and according to [45] and [46] as a consequence may occur: 1) reduced prostaglandin (PG) synthesis of series 2; 2) reduction of thromboxane A2 (TXA2), which acts by aggregating platelets and causes vasoconstriction; 3) reduced production of leukotrienes B4, which induces inflammation; 4) increased of thromboxane A3 (TXA3), which are weak as platelet aggregators and as vasoconstrictors; 5) higher synthesis of prostaglandin I3 (PGI3) without reducing PGI2, which act as active vasodilators and inhibitors of platelet aggregation; 6) higher concentrations of LT of type B5, which is a chemotactic agent and a weak inducer of inflammation.

Lymphocytes become active and initiate cytokine secretion stimulating the immune response, although their proliferation may be impaired due to γ-linolenic acid or EPA [47,48]. Caughey et al. [49] suggest a reduction in the function of immune cells after supplementing humans with a high concentration of n-3 PUFA (linseed oil) due to reduced synthesis of interleukin-1 and tumor necrosis factor (TNF) by blood monocytes.

Ingestion of PUFA leads to their distribution to virtually every cell in the body, resulting in effects on membrane composition and function, cellular signaling, and regulation of gene expression through various transcription factors. Although the results obtained for gilt in growing were not expressive in the current study, more studies are necessary to clarify how the consumption of cow’s milk enriched naturally with PUFA can influence the development of animals e humans.

In conclusion, gilt that received the cow’s milk enriched naturally with PUFA n-3 or n-6 had similar growth performance, but the blood fatty acid profile and hematological parameters were altered without impairing the health of gilts. This study provided insight concerning how food enriched can modulate the serum lipid profile and cells of the defense system, promoting health benefits.

## ABBREVIATIONS

ADFI: average daily feed intake
ADG: average daily gain
EPA: eicosapentaenoic acid
FCR: feed conversion ratio
HDL: high-density lipoproteins
LTs: leukotrienes
LDL: low-density lipoproteins
MCH: mean corpuscular hemoglobin
PUFA: polyunsaturated fatty acids
RBC: red blood cells
SCD: stearoyl-CoA desaturase
SFA: saturated fatty acids
USFA: unsaturated fatty acids
VLDL: very-low-density lipoprotein.

## ACKNOWLEDGEMENTS

The authors would like to thank the São Paulo Research Foundation (FAPESP) by the financial support (Grant n° 2015/19393-8) and Coordenação de Aperfeiçoamento de Pessoal de Nível Superior – Brasil (CAPES) – Finance Code 001.

The authors’ contributions are as follows: A. S. N. designed the study and was the principal investigator who supervised all aspects of the study. A. S. N. and A. F. C. d. A. conceptualized the study. L. G. R. conducted the animal study and wrote the manuscript as part of her postgraduate studies. T. H. S. realized the statistical analysis. S. M. M. K. M. critically reviewed the manuscript. L. G. R., T. H. S., S. M. M. K. M., M. S. V. S. and P. L. T. did data curation. All the authors read, edited, and approved the final manuscript.

The authors declared that there are no conflicts of interest.

